# Comparison of alternative solvents for *in situ* extraction of hydrocarbons from the colonial green alga *Botryococcus braunii* race B (Showa)

**DOI:** 10.1101/2023.10.23.563540

**Authors:** Takehiro A. Ozawa-Uyeda, Sebastian Overmans, Bárbara Bastos de Freitas, Edmundo Lozoya-Gloria, Kyle J. Lauersen

**Affiliations:** Genetic Engineering Department, CINVESTAV-IPN Irapuato Unit, C.P. 36824 Irapuato, Gto., Mexico; Bioengineering Program, Biological and Environmental Sciences and Engineering Division, King Abdullah University of Science and Technology (KAUST), Thuwal 23955-6900, Saudi Arabia

**Keywords:** *Botryococcus braunii*, microalgae, *in situ* extraction, hydrocarbons, green solvents

## Abstract

The colony-forming, green microalga *Botryococcus braunii* secretes petroleum-like hydrocarbons, which enables the non-destructive continuous *in situ* extraction, ‘milking’, of these extracellular products during culture growth without cell lysis. This work compares the suitability of 15 different solvents, including alkanes, halogenated solvents, and green solvents, for *in situ* extraction of *B. braunii* race B (Showa strain) hydrocarbons after acclimation to moderate salinity stress. After 24 h of extraction, bio-based terpene ‘green’ solvents such as ɣ-terpinene showed the highest hydrocarbon recovery, around 10-fold greater than with conventional alkane solvents. Brominated alkanes and liquid perfluorocarbons (FCs) formed a lower phase to algal cultures rather than an upper phase as with other solvents, but only bromodecane effectively captured extracellular hydrocarbons similar to conventional alkane solvents. However, bromodecane and all green solvents were too toxic for two-phase continuous culture contact hydrocarbon milking, leading to 33–100% chlorophyll content loss. To overcome the biologically adverse effects of these solvents with suitable hydrocarbon recovery, future research should focus on their application in short-term extraction period ‘milking’ systems to minimize algal-solvent contact and enable continuous extraction.

Graphical Abstract

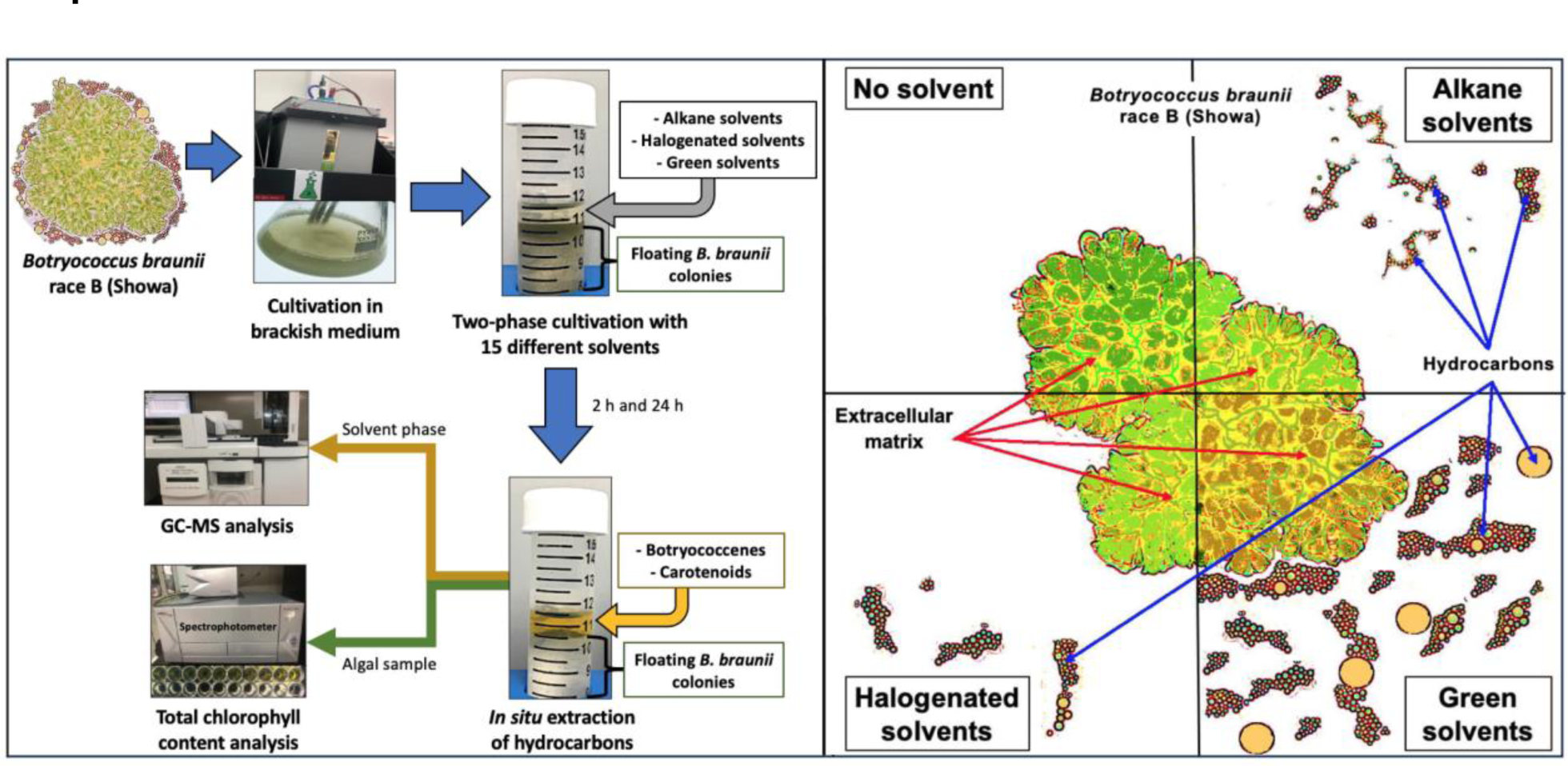

## Introduction

The colony-forming green microalga *Botryococcus braunii* race B produces and secretes large amounts of liquid triterpene hydrocarbons, botryococcenes, during its exponential growth phase, which are stored in its extracellular matrix (ECM) (Largeau *et al*., 1980; Suzuki *et al*., 2013). Botryococcenes can be used to produce petroleum-equivalent fuels suitable for combustion engines (Jin *et al*., 2016; Lozoya-Gloria *et al*., 2019). In contrast to most other oleaginous microalgae, which accumulate oils within the cell, extracellular *B. braunii* hydrocarbons can be harvested without requiring cell lysis (Suzuki *et al*., 2013; Ennaceri *et al*., 2023). This trait makes it a suitable microalga for the process of non-destructive *in situ* extraction, termed ‘milking’, of lipophilic compounds. Milking can eliminate the steps of algal biomass downstream processing: harvesting, dewatering and cell disruption (Jackson *et al*., 2017; Kleinert and Griehl 2022; Ennaceri *et al*., 2023).

*B. braunii* typically exhibits slow growth rates compared to other photosynthetic microbes, which has limited its broader biotechnological exploitation (Lozoya-Gloria *et al*., 2019). Additionally, the colony surface of *B. braunii* race B is covered with an ultrastructure called a retaining wall, which is decorated with a fibrillar sheath composed mainly of arabinose-galactose polysaccharides (Weiss *et al*., 2012). This outer barrier hinders the entry of many organic solvents, which limits milking efficiencies (Furuhashi *et al*., 2016b). One pretreatment that was found to improve the extraction rate of botryococcenes is cultivation of the alga in brackish medium (Furuhashi *et al*., 2013; 2016a). It was proposed that increased salinity in the culture medium decreased the production or integrity of the polysaccharides that block non-polar solvents and increases colony size and flotation (Furuhashi *et al*., 2016a; 2016b).

Extraction of hydrocarbons from *B. braunii* has been tested with different types of solvents, including traditional alkanes solvents such as hexane (C6) and heptane (C7), partially polar solvents such as 1-octanol, and alternatives solvents such dimethyl ether (DME) and switchable-polarity solvents (SPS) (Frenz *et al*., 1989; Samori *et al*., 2010; Kanda *et al*., 2013). Short chain alkanes are reported to have high botryococcene extraction efficiencies but low biocompatibility, resulting in rapid loss of cell viability after long contact times (Kleinert and Griehl, 2021). Dodecane (C12) is biocompatible with *B. braunii* cultures but has reduced hydrocarbon extractability compared to hexane (Mehta *et al*., 2019). Alkane solvents have significant drawbacks that hinder scaling their application to larger culture volumes like flammability and emulsion formation at the culture-solvent interface (Lauersen, 2019).

It was recently reported that halogenated perfluorocarbon (FC) liquids can enable milking of sesqui- and di-terpenoids from engineered algal cultures (Overmans and Lauersen, 2022). These have not yet been tested on *B. braunii*. Green solvents are another class of chemicals that are bio-based in origin and have gained popularity in process chemistry due to their biological origins and/or biodegradability (Kumar *et al*., 2017; De Jesus and Maciel Filho, 2020). Monoterpenes derived from essential oils such as D-limonene, α-pinene, and p-cymene are considered ‘green’ solvents because they are obtained from renewable feedstocks, like plants and crop wastes (Kumar *et al*., 2017). Lipid extraction of microalgae and oilseeds using terpene green solvents was shown to have equal or superior quality and yields than with hexane (De Jesus and Maciel Filho, 2020). In this study, we evaluated the suitability of halogenated and green solvents as alternatives to conventional alkanes for botryococcene extraction from *B. braunii* race B (Showa strain) cultures grown in brackish media.

## Materials and Methods

### Reagents

The solvents tested in this work were grouped into straight-chain alkanes, halogenated solvents, and green solvents. Alkanes included n-hexane, n-decane, n-dodecane (≥99%), and n-pentadecane, which were obtained from Acros Organics (Geel, Belgium), VWR International (Fontenay-sous-Bois, France), Sigma-Aldrich (St. Louis, USA) and Alfa Aesar (Heysham, England), respectively. Halogenated solvents included brominated alkanes (bromododecane and bromododecane) and chlorinated alkane (chlorodecane), acquired from Sigma Aldrich (St. Louis, USA). Additionally, two perfluorocarbon liquids, hereafter referred to as FCs, were used in this work: the perfluorinated ether FC-770 and the perfluorinated amine FC-3283, obtained from Fluorochem (Glossop, UK) and Acros Organics (Geel, Belgium), respectively. The green solvents included three monoterpenes (para-cymene (99%), gamma-terpinene, and alpha-pinene) and three alcohols (1-heptanol, nerol, and nopol), all purchased from Sigma-Aldrich (Taufkirchen, Germany). According to GlaxoSmithKline’s (GSK) green solvents guide for assessing green credentials of solvents, 1-heptanol scored highly compared to other solvents, such as para-cymene and n-hexane (Alder *et al*., 2016). For chlorophyll quantification, ethanol (96% vol) and methanol were purchased from VWR International (Fontenay-sous-Bois, France).

### Microalgal cultures and growth conditions

Batch cultures of *Botryococcus braunii* race B, Showa strain (Nonomura, 1988), were grown in 1-L Erlenmeyer flasks in Algem photobioreactors (Algenuity©, UK) at 25 °C under 130 μmol photons m^−2^ s^−1^ with a 12h:12h light:dark cycle and sparged with a 7% CO_2_ in air mixture. The gas mix was supplied periodically to cultures at 25 cc min^−1^ according to the pH control level set on the photobioreactor, with the online pH measured continuously using a calibrated pH probe (Broadley James, UK) to maintain a constant pH of 7.5 in the culture media. *B. braunii* was cultured in biological triplicates (n = 3) in three different media: modified Chu-13 medium as the standard freshwater medium based on the recipe in (Furuhashi *et al*., 2013), modified Chu-13 medium with 37.5 mM NaCl, and modified Chu-13 medium with 5% Red Sea water (RSW).

To avoid a severe algal growth inhibition due to relatively high salinity (e.g. 150 mM NaCl in 1/4 seawater medium) as previously reported (Furuhashi *et al*., 2013), the artificial seawater media were reduced to 3–4 g L^-1^ brackish salinity medium (Furuhashi *et al*., 2016a; 2022). Culturing *B. braunii* race B (Showa strain) and *B. braunii* race A (Yamanaka strain) in brackish media with 3 and 4 g L^-1^ salinity, respectively, improved their hydrocarbon extractability without inhibiting algal growth compared to the algae cultured in Chu13 freshwater medium (Furuhashi *et al*., 2016a; 2022). Instead of assuming that all the salt in the seawater is NaCl, the content of NaCl accounted for 57.6% of dissolved salt in the artificial seawater medium used by Furuhashi *et al*. (2016a). Therefore, 3 and 4 g L^-1^ salinity medium contained 30 and 40 mM NaCl, respectively. Brackish medium was prepared by mixing filtered (0.2 μm) and autoclaved RSW with modified Chu-13 medium. The concentration of total dissolved solids in RSW was previously determined to be around 750 mM NaCl (Abdel-Aal *et al*., 2015). To get a final NaCl concentration within the range of 30–40 mM, similar to the most suitable concentrations used by Furuhashi *et al*. (2016a; 2022), Chu-13 medium was adjusted to a final concentration of 5% RSW (37.5 mM NaCl). All cultures were grown in the Algem photobioreactors with a working volume of 400 mL for 18 days before being recovered for further analyses.

### Determination of growth parameters

Final algal biomass was determined gravimetrically by obtaining a pellet of 5 mL algal culture after adding an equal volume of 96% ethanol into a pre-weighed glass tube, mixed manually several times, and centrifuged at 3,000 × *g* for 1 min to sediment all algal cells. The addition of ethanol helps to precipitate the floating layer of algal colonies, but the incubation time should be as short as possible to minimize extraction of undesired pigments. After discarding the ethanol, the remaining algal pellet was dried in an oven at 110 °C for 24 h and cooled for 30 min before weighing. The dry cell weight (DCW; in mg) of the microalgae was obtained by subtracting the dry weight of the empty glass tube from the loaded glass tube. The specific growth rate was calculated using the following equation:

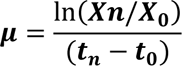

where, *µ* is the specific growth rate (day^-1^), *Xn* is the final biomass concentration (g/L) at time *t_n_* (days), and *X_0_* is the initial biomass concentration (g/L) at time *t_0_* (days).

Total chlorophyll content was obtained by adding 1 mL 96% ethanol to 1 mL culture samples, mixed by inversion several times, and centrifuged at 11,000 × *g* for 1 min to sediment all suspended algal cells. After discarding the ethanol, the pellet was resuspended in 1 mL 100% methanol and incubated for 30 min at 45 °C in the dark to extract chlorophyll. The cell debris was pelleted by centrifugation at 11,000 × *g* for 1 min while the supernatant was used to measure the absorbance at 666, 653, and 470 nm with a spectrophotometer (TECAN Infinite M1000 PRO). Total chlorophyll content was calculated according to Wellburn (1994). The values of total chlorophyll content were then expressed on a dry weight basis (mg chlorophyll/ g DCW) using the DCW of the algal samples.

### Determination of hydrocarbon content

Algal pellet samples were collected by centrifugation as described above, and DCW was determined. Hydrocarbons were extracted from the dried algal sample by adding 5 mL of n-hexane and incubating for 30 min at room temperature with manual shaking at 10 min intervals. The extraction solvent containing botryococcene hydrocarbons, which turned yellowish-orange due to the carotenoids found within the ECM (Eroglu and Melis, 2010), was separated from the algal residue by decantation and collected in a separate glass flask. This process was repeated until the solvent phase became colorless. All n-hexane extracts were combined and evaporated in an oven at 110 °C overnight. Hydrocarbon amounts were calculated gravimetrically and expressed as a percentage of biomass DCW. For positive identification of botryococcenes in samples, the extracts were resuspended in 1 mL of n-hexane or n-dodecane, and then transferred to gas chromatography (GC) vials (n = 3) and stored at −20 °C until further analysis as described below.

### Preliminary screening of solvents for two-phase ‘milking’ of *B. braunii* race B in low salinity culture medium

Two-phase living extractions (i.e., ‘milking’) of extracellular hydrocarbons were performed with alkanes, halogenated-, and green solvents by gently adding 250 µL of each respective solvent (25% v/v) onto 750 µL liquid cultures of *B. braunii* that were previously grown in brackish medium for 18 days. All two-phase cultures were grown for 24 h in duplicates in 2-mL microcentrifuge tubes on an orbital shaker at 120 rpm with a 12h:12h light:dark cycle at room temperature. The solvent layer was then collected following centrifugation at 4,500 × *g* for 5 min to obtain two separate layers. From each sample, 150 μL of the solvent fraction was transferred into individual GC vials and stored at −20 °C until further analysis, as described below.

### Alternative solvents for *in situ* extraction of hydrocarbons produced by *B. braunii* race B

The two most efficient extraction solvents of each group were selected from the initial screening. Two-phase living extractions of extracellular hydrocarbons were performed by gently adding 1 mL of each respective solvent onto 9 mL liquid cultures of *B. braunii* (10% v/v) that were previously grown in a brackish medium for 18 days. The two-phase cultures were cultivated for 24 h in triplicates in 50-mL Erlenmeyer flasks on an orbital shaker at 120 rpm with a 12h:12h light:dark cycle at room temperature. The solvent and algal samples were obtained at two different time points: 2 h and 24 h of cultivation. At each time point, the solvent layer was collected after centrifugation at 4,500 × *g* for 5 min. From each sample, 150 μL of the solvent fraction was transferred into separate GC vials and stored at −20 °C until further analysis.

### Evaluation of hydrocarbon recovery rate

Hydrocarbon extractability from *B. braunii* culture was expressed as the hydrocarbon recovery rate, defined as the percentage of botryococcene hydrocarbons extracted from the wet algal samples using different types of solvents compared to the amount extracted from the dried algal sample using hexane as described above. The extracted botryococcene content was determined from the sum of two main peak areas detected at retention times of 9.7 min. and 10.2 min., per mg of dry algal cell weight.

### Gas chromatography

All solvent extracts were analyzed using a gas chromatograph equipped with a mass spectrometer and a flame ionization detector (GC-MS-FID) with modifications to a previously described protocol (Thapa *et al*., 2016). Briefly, the GC-MS-FID analyses were performed using an Agilent 7890A gas chromatograph connected to a 5975C inert MSD with a triple-axis detector. The system was equipped with a DB-5MS column (30 m × 0.25 mm i.d., 0.25 μm film thickness) (Agilent J&W, USA). The temperatures were set to: injector (260 °C), interface (250 °C), and ion source (200 °C). One µL of liquid solvent sample was injected in splitless mode with an autosampler (Model G4513A, Agilent). Column flow was kept constant at 2.5 mL min^−1^ with He as a carrier gas. The initial GC oven temperature of 220 °C was held for 1 min, then raised to 280 °C at a rate of 5 °C min^−1^, followed by 2 °C min^−1^ to 300 °C and held for 5 min. Mass spectra were recorded after a 3 min solvent delay using a scanning range of 50–750 m/z at 20 scans s^−1^.

Gas chromatograms were manually reviewed for quality control before botryococcene GC peak areas were integrated using MassHunter Workstation software version B.08.00 (Agilent Technologies, USA). The NIST library (National Institute of Standards and Technology, Gaithersburg, MD, USA) was used to identify botryococcenes.

### Statistical analysis

Differences between the mean values obtained from each assay were assessed through one-way analysis of variance (ANOVA) followed by Tukey’s HSD post-hoc test with a confidence level of 95%. All statistical analyses were performed using R software v3.4.3 (https://www.r-project.org/) through the RStudio interface.

## Results and Discussion

### *B. braunii* growth and hydrocarbon extractability in different media

After 18 days of cultivation, there were no significant differences in specific growth rates of *B. braunii* cultured in modified Chu13 medium (0.077 ± 0.002 day ^−1^; mean ± SEM) and in 5% RSW (0.064 ± 0.007 day ^−1^), but the algae cultured in 37.5 mM NaCl (0.050 ± 0.005 day ^−1^) showed a significant decrease (*p* < 0.01) in the specific growth rate compared to the other treatments (Fig. 1A). *B. braunii* cultured in 5% RSW and in 37.5 mM NaCl had significant decreases in chlorophyll content (3.6 ± 0.40 mg g^−1^ DCW and 3.4 ± 0.28 mg g^−1^ DCW, respectively) and appeared less healthy (chlorotic phenotype) compared to algae cultured in non-saline medium (5.3 ± 0.13 mg g^−1^ DCW) (Fig. 1B).

**Figure 1.**
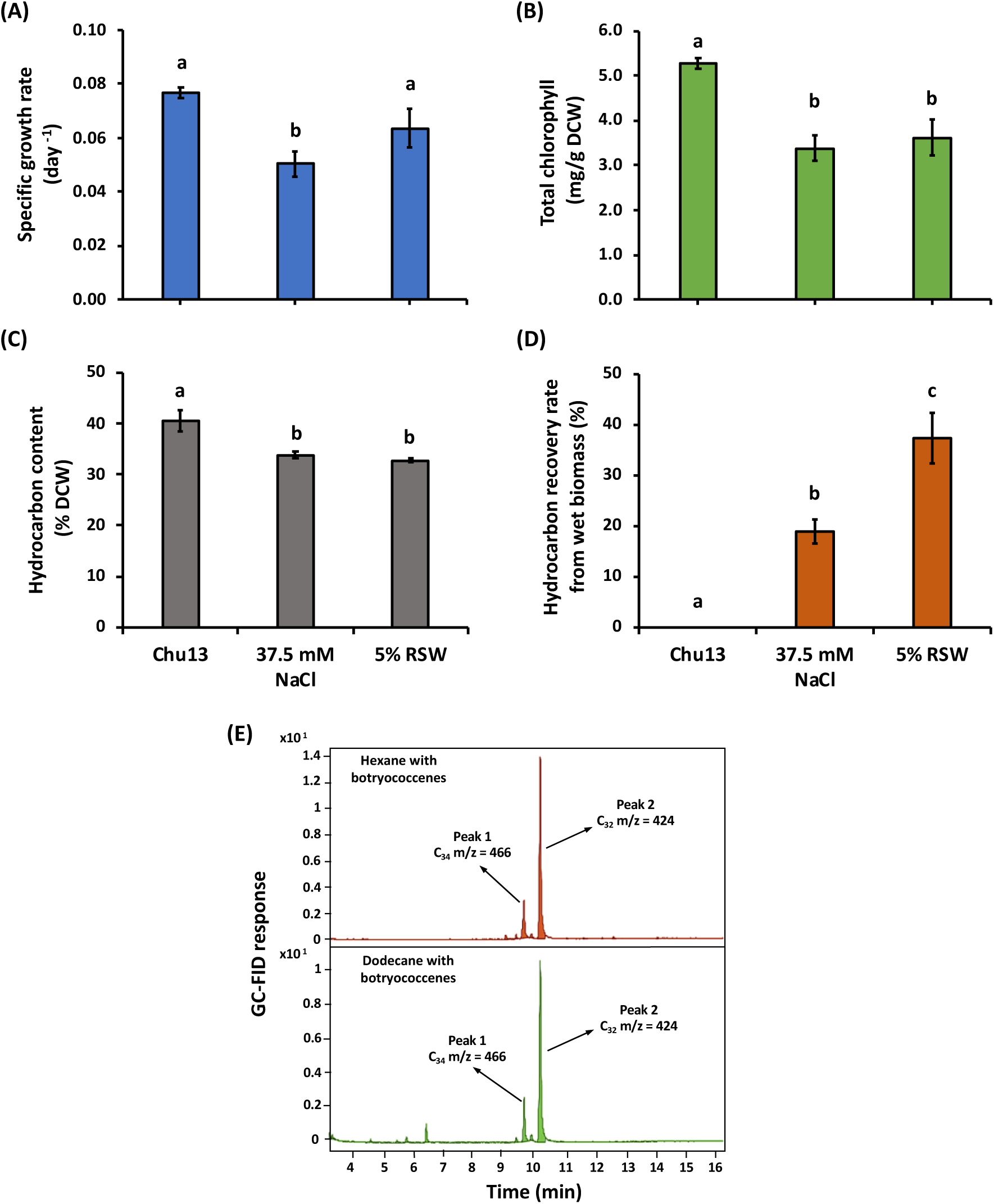
Analysis of *B. braunii* (Showa strain) cultured for 18 days in three different media. (**A**) Specific growth rate, (**B**) total chlorophyll content, (**C**) extracellular hydrocarbon content, and (**D**) hydrocarbon recovery rate of *B. braunii* race B cells grown in modified Chu-13 medium (Chu13), modified Chu-13 medium with 37.5 mM NaCl (37.5 mM NaCl), and modified Chu-13 medium with 5% Red Sea water (5% RSW). Recovered hydrocarbons from dry algae by hexane extraction accounted for a 100% hydrocarbon recovery rate. (**E**) Representative GC-FID chromatograms of botryococcenes extracted with commonly used alkane solvents obtained from wet *B. braunii* algal samples. Error bars represent the standard error of the mean of three biological replicates (SEM; n = 3). Statistical significance was determined by one-way ANOVA with Tukey’s HSD post-hoc test (p < 0.05). Within each panel, different letters above bars indicate instances where means were significantly different (*p* < 0.05).

The hydrocarbon content in dry biomass of *B. braunii* cultured in the brackish medium was 33 ± 0.5% DCW compared to 40 ± 2.1% DCW in modified Chu13 medium (Fig. 1C). However, the hydrocarbon recovery rate from suspended *B. braunii* cells in 5% RSW (37.3 ± 5.1%) was 2-fold greater than in the suspended algae cultured in 37.5 mM NaCl conditions (18.8 ± 2.4%), while the hydrocarbon extractability in non-saline medium was non-existent (Fig. 1D). The presence of C_34_ and C_32_ botryococcenes from suspended *B. braunii* cells cultured in 5% RSW was confirmed by the appearance of two distinct peaks (9.7 min and 10.2 min, respectively) with appropriate mass spectra in GC-MS chromatograms (Fig. 1E). These results indicate that culturing *B. braunii* in modified Chu-13 medium with 5% RSW is possible without strongly reducing its growth and encourages the production of extracellular botryococcenes. This is in line with previous reports (Furuhashi *et al*., 2013; 2016; 2022), where *B. braunii* cultivation in brackish medium reduced the cultures’ growth rate but improved hydrocarbon extractability from algal biomass compared to cultivation in non-saline medium.

### Suitability screening of solvents for two-phase cultivation of *B. braunii* race B in brackish culture medium

A recent study has shown that the Showa strain is one of the most promising for *in situ* extraction because of its high solvent resistance, hydrocarbon extractability, and lipid productivity (Kleinert and Griehl 2021). Our results showed that modified Chu-13 medium with 5% RSW was the best for growing and improving botryococcene recovery rate around cultured *B. braunii* Showa (Fig. 1). Because *B. braunii* hydrocarbons are mainly (90–95%) stored in the ECM, most of them can be extracted by thermal pretreatment (i.e. drying or heating of wet algae) or culturing the algae in brackish medium before soaking the algae in a non-polar organic solvent such as hexane that does not permeate the cells (Largeau *et al*., 1980; Suzuki *et al*., 2013; Furuhashi *et al*., 2016b). Hydrocarbons obtained in this way are typically referred to as extracellular hydrocarbons. After recovering the extracellular hydrocarbons, a small amount of intracellular hydrocarbons can also be extracted with a mixture of chloroform and methanol that can reach inside the cells (Largeau *et al*., 1980). For the initial solvent screening, cultures grown in 5% RSW were treated with 14 different non-polar solvents at 25% (v/v), whose physicochemical properties are shown in Table 1.

**Table 1.**
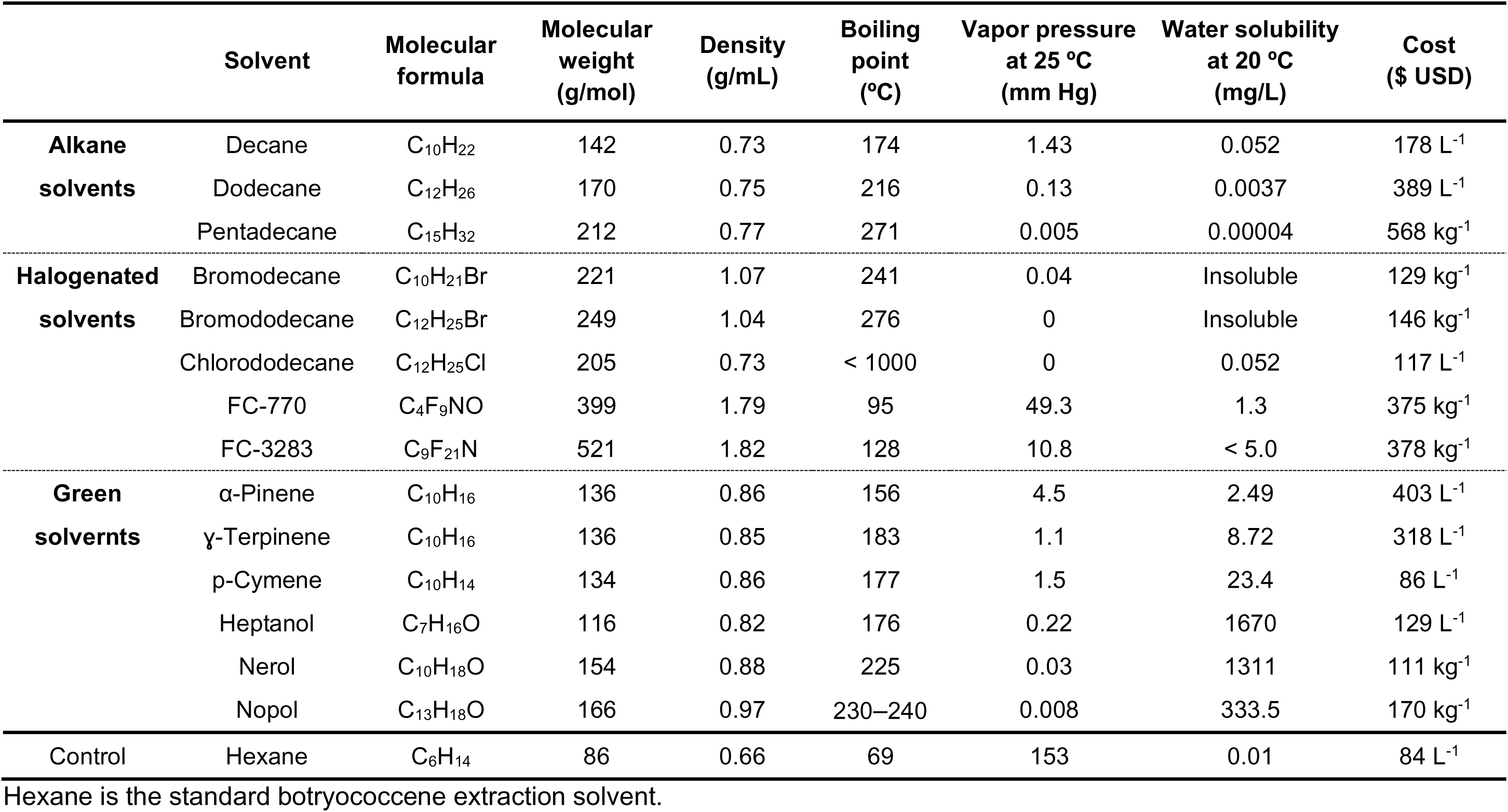
Physico-chemical properties of the solvents used in this work.

After 2 and 24 h extraction, the solvents exhibited a highly variable performance in extracting botryococcene hydrocarbons from the algal cells (Fig. 2A). This was visually indicated by the co-extraction of extracellular carotenoids that had previously been reported to be associated with the botryococcene fraction from the Showa strain (Eroglu and Melis, 2010). Most of the solvents showed yellowish coloration at different levels in the upper phase (Fig. 2A). Brominated alkanes and FCs formed a lower phase (Fig. 2A). Alkane solvents showed a similar but relatively poor hydrocarbon recovery rates (Fig. 2B). Halogenated solvents were worse than alkanes except for bromodecane, which was the best of all previously tested solvents (Fig. 2B). However, the hydrocarbon recovery rate with green solvents (e.g. 34.2 ± 0.5% with α-pinene) was found to be up to ∼10-fold and ∼3.8-fold greater than with dodecane (3.3 ± 0.1%) and bromodecane (8.9 ± 1.2%), respectively (Fig. 2B). The exception was nopol which did not extract botryococcenes (Fig. 2B). This indicates that green solvents, such as ɣ-terpinene and α-pinene, were more efficient at extracting botryococcenes *in situ* compared to conventional alkane solvents and their halogen derivatives (Fig. 2B).

**Figure 2.**
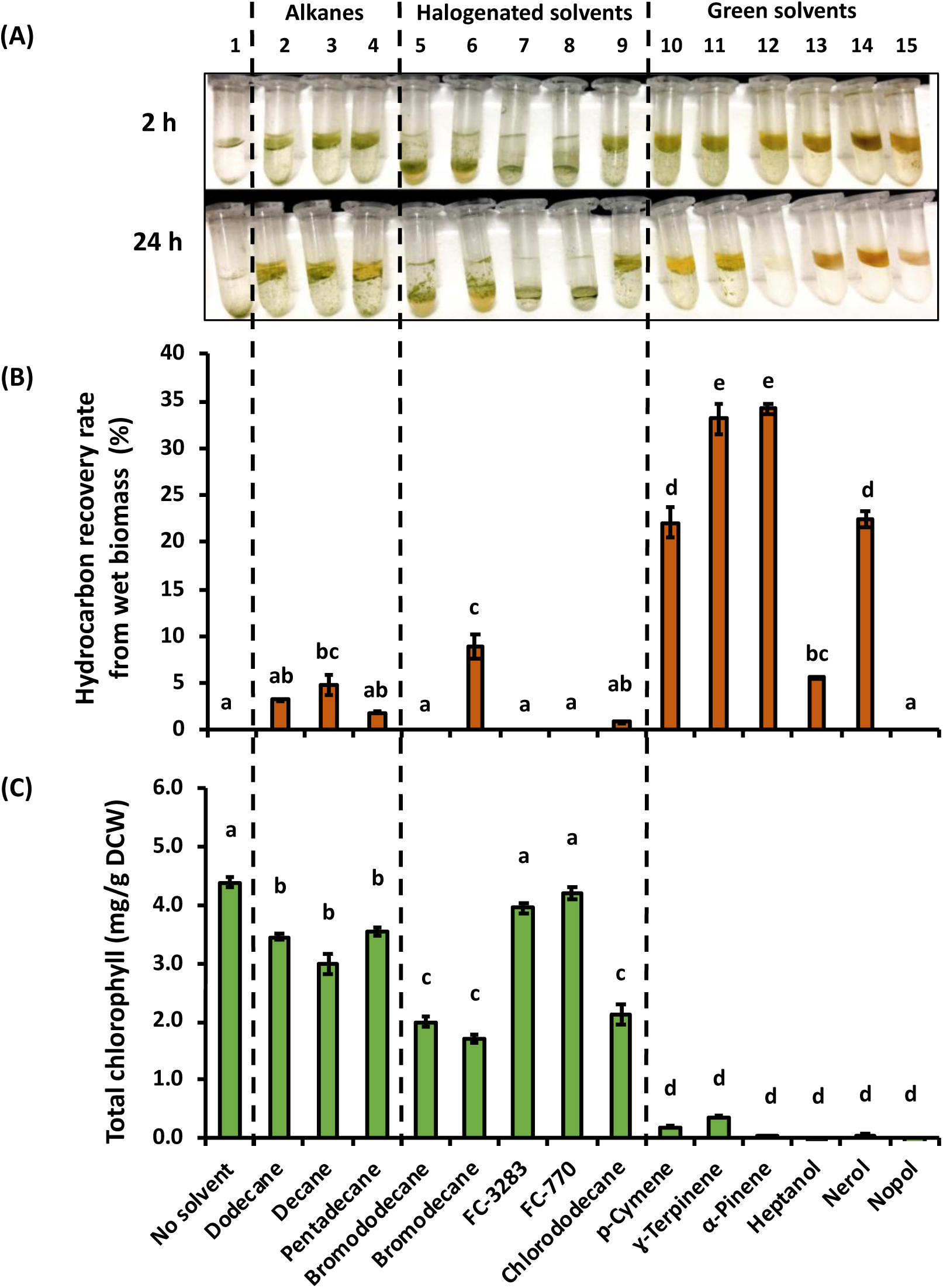
Extraction and toxicity properties of all different solvents. (**A**) *B. braunii* race B cells grown in culture medium with 5% RSW after 2 h and 24 h extraction with 25% (v/v) of solvents. (**B**) Hydrocarbon recovery rates after 24 h of two-phase living *B. braunii* extraction cultures with different solvents. Recovered hydrocarbons from dry algae by hexane extraction accounted for a 100% hydrocarbon recovery rate.(**C**) Total chlorophyll content in *B. braunii* cultured in 5% RSW medium after 24 h extraction with each solvent. Error bars represent the standard error of the mean of two biological replicates (SEM; n = 2). Statistical significance was determined by one-way ANOVA with post-hoc Tukey’s HSD test. Letters above the bars indicate significantly different mean concentrations (*p* < 0.05).

Although both α-pinene and ɣ-terpinene yielded the highest extraction efficiency of botryococcenes in this screening (Fig. 2B), we observed a severe degradation of chlorophyll in all green solvents extracts after 24 h of incubation compared to alkanes and halogenated solvents (Fig. 2C). The total chlorophyll content in *B. braunii* after two-phase cultivation with solvents was measured to evaluate solvent biocompatibility. The addition of solvents to *B. braunii* led to differing extents of chlorosis. Of the different two-phase cultivations, algal cultures with alkanes showed a slight decrease in the chlorophyll content compared to cultivations without solvent (Fig. 2C). Algae with brominated alkanes and chlorododecane demonstrated a stronger reduction in chlorophyll content (1.7–2.1 mg g^−1^ DCW) than those with straight-chain alkanes (3.0–3.6 mg g^−1^ DCW) while cultures with FCs had chlorophyll contents (3.9–4.2 mg g^−1^ DCW) similar to the no-solvent treatment (4.4 ± 0.08 mg g^−1^ DCW) (Fig. 2C). Green solvents were the most toxic solvents, particularly the alcohols, leading to a drastic decrease in chlorophyll content to around 0–0.4 mg g^−1^ DCW (Fig. 2C).

Our results indicate that green solvents are too toxic for two-phase cultivations, unlike longer chain alkanes and halogenated solvents. A previous study reported similar results after comparing the effects of decane, its alcohol derivative decanol, and the green solvent D-limonene, where decanol was the most toxic solvent, followed by D-limonene and decane, (Concha *et al*., 2018). Here, brominated alkanes and FCs formed a lower phase or ‘underlay’ with algal cultures, unlike other solvents that formed a distinct layer on top of the culture (Fig. 2A). Of the heavy solvents, only bromodecane could capture hydrocarbons from wet algal samples (Fig. 2B). In contrast, previous work reported that FCs could replace the use of dodecane to extract the sesqui- and diterpenoids produced by engineered green microalgae (Overmans and Lauersen, 2022).

### Selected solvents for *in situ* extraction of hydrocarbons produced by *B. braunii* race B

We evaluated *in situ* hydrocarbon extraction in larger culture volumes at two incubation periods, 2 h and 24 h, with six solvents: hexane, dodecane, bromodecane, bromododecane, p-cymene and ɣ-terpinene. *B. braunii* cultured in 5% RSW medium were gently overlaid with each solvent at 10% (v/v), which is the standard culture:solvent ratio used to capture terpenoids or hydrocarbons from engineered algae (Overmans and Lauersen, 2022; Yunus *et al*., 2022). Regardless of solvent density, we observed the formations of emulsions in cultures with all solvents after shaking the mixtures at 120 rpm even after 2 h (Fig. 3A). This formation was previously identified as one technical limitation for isoprenoid milking with hydrophobic solvents on algal culture (Lauersen, 2019). After 2 h of extraction, the alkane solvents dodecane and hexane showed a yellow coloration, indicating hydrocarbon extraction, while the aqueous lower phase contained algae and culture medium. After 24 h of extraction, algal cultures were only green in dodecane, which also exhibited a more intense yellow color accumulation than hexane, where cells were chlorotic or dead (Fig. 3A).

**Figure 3.**
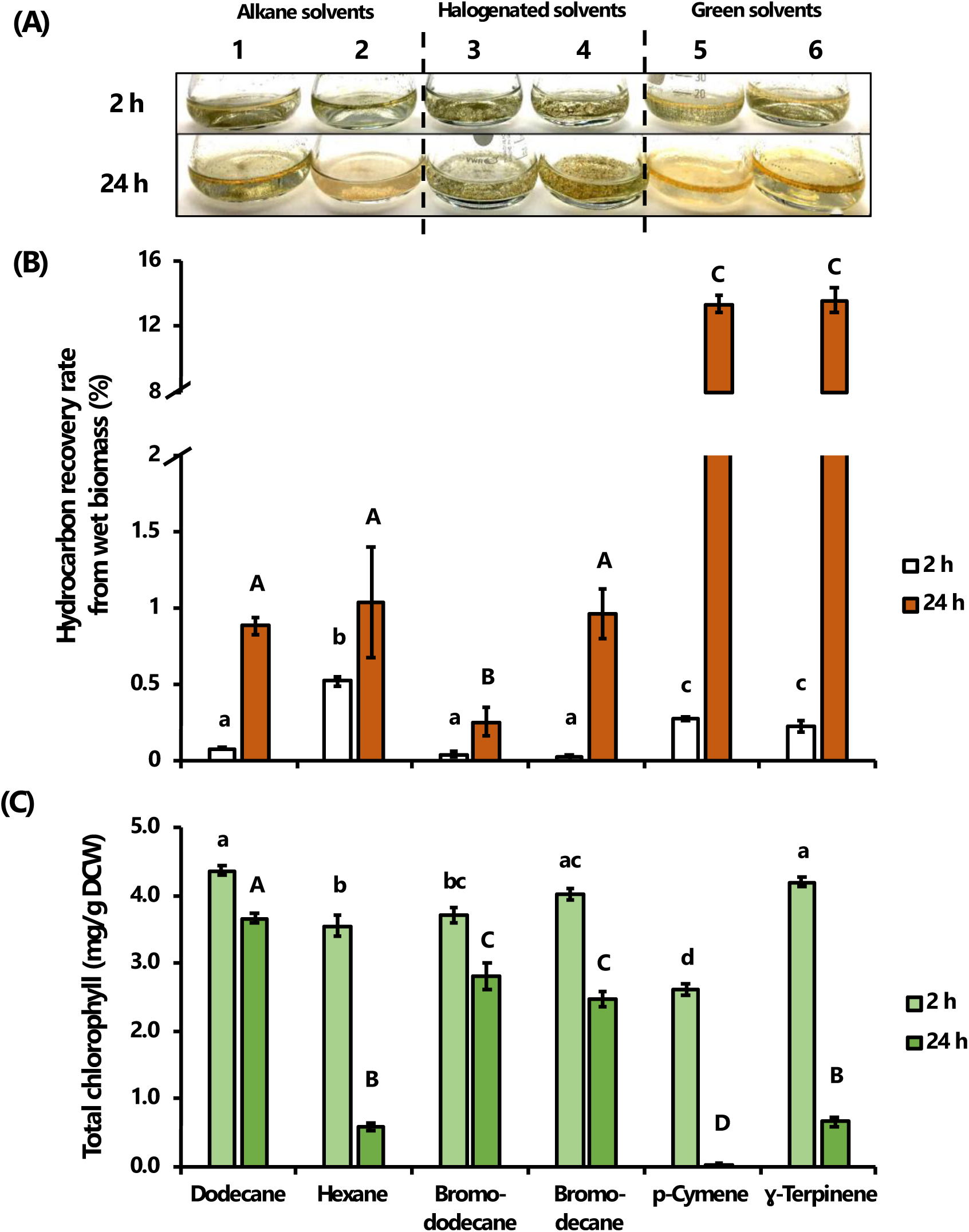
Extraction and toxicity properties of selected solvents. (**A**) *B. braunii* cells grown in culture medium with 5% RSW after a contact time of 2 h and 24 h with 10% (v/v) of either (1) hexane, (2) dodecane, (3) bromododecane, (4) bromodecane, (5) p-cymene, or (6) ɣ-terpinene. (**B**) Hydrocarbon recovery rates after 2 h and 24 h of two-phase living *B. braunii* extraction cultures with solvents. Recovered hydrocarbons from dry algae by hexane extraction accounted for a 100% hydrocarbon recovery rate. (**C**) Total chlorophyll content of *B. braunii* cultured in medium with 5% RSW after 2 h and 24 h extraction time with solvents. Error bars represent the standard error of the mean of three biological replicates (SEM; n = 3). Statistical significance was determined by one-way ANOVA with post-hoc Tukey’s HSD test. Lower- and upper-case letters above the bars indicate the post hoc results of the 2 h and 24 h data sets, respectively. Means were considered significantly different at level *p* < 0.05.

Brominated solvents formed a lower phase to the aqueous algal culture where colonies clumped, but no hydrocarbon extraction occurred from either solvent. Terpene green solvents were found to extract the hydrocarbons, with aggregation of colonies visible at 2 h and severe chlorophyll degradation at 24 h of extraction (Fig. 3A). After a 2 h contact time between solvents and the 5% RSW-acclimated *B. braunii* cells, we detected the highest hydrocarbon content in hexane, followed by p-cymene and ɣ-terpinene (Fig. 3B). The hydrocarbon recovery rate in extractions with dodecane and brominated alkanes were found to be much higher after 24 h than after 2 h (Fig. 3B). Bromodecane showed a similar hydrocarbon extractability compared to the two alkanes after 24 h of incubation. Even though the hydrocarbon recovery rate with bromodecane resulted in a 3.8-fold increase compared to bromododecane (Fig. 3B), both of the tested brominated alkanes caused a similar decrease in chlorophyll content compared to dodecane from 3.7 ± 0.08 mg g^−1^ DCW to ∼2.6 mg g^−1^ DCW (Fig. 3C). *B. braunii* race B cells cultured with hexane and p-cymene showed a fast decrease in chlorophyll content (3.56 ± 0.16 mg g^−1^ DCW and 2.62 ± 0.08 mg g^−1^ DCW, respectively), whereas algal cells with ɣ-terpinene (4.21 ± 0.06 mg g^−1^ DCW) showed a similar chlorophyll content to cells with dodecane after 2 h of incubation (4.37 ± 0.07 mg g^−1^ DCW) (Fig. 3A and 3C). Longer contact time (24 h) with ɣ-terpinene was detrimental to algal cells (0.66 ± 0.06 mg g^−1^ DCW), as could be seen in the hexane treatment (Fig. 3A and 3C). This result indicates that the bromodecane underlay was amenable to the extraction of extracellular hydrocarbons and carotenoids from *B. braunii* cells, but it was more aggressive as a two-phase culture system than the conventional dodecane overlay.

After 24 h of extraction, *B. braunii* cultured in brackish medium with 10% (v/v) of solvents showed lower hydrocarbon recovery rates compared to the algae cultured with 25% (v/v) of solvents (Fig. 2B and 3B). Increasing culture salinity to 10% RSW (75 mM NaCl) could improve the hydrocarbon extraction efficiency from suspended algal cells without requiring a high solvent content. It was shown that the hydrocarbon recovery rate from *B. braunii* race B cultured in brackish medium with a salinity of 70 mM NaCl for 18 days was nearly equal to that from the freeze-dried algae (100% hydrocarbon recovery rate), but the biomass production was reduced (Furuhashi *et al*., 2016a). Another strategy could focus on milking hydrocarbons from wet algae in their early growth phase after each subculture into fresh brackish medium. For example, it was reported that *B. braunii* race B cultured in brackish media with a salinity of 30 and 50 mM NaCl for 18 days showed a hydrocarbon recovery rate of around 12% and 36% of dry heptane extraction, respectively. However, the algae on day 24, which was 6 days after subcultivation in brackish media with a salinity of 30 and 50 mM NaCl, showed an increase in the hydrocarbon recovery rate of around 35% and 90% of dry heptane extraction, respectively (Furuhashi *et al*., 2016a).

The highest hydrocarbon recovery rates were detected in p-cymene (13.3 ± 0.5%) and ɣ-terpinene (13.6 ± 0.8%) extracts after 24 h of incubation, both showing hydrocarbon extractabilities 13- and 15-fold greater than hexane (1.0 ± 0.4%) and dodecane (0.9 ± 0.1%), respectively (Fig. 3B). As far as we know, this is the first report of using γ-terpinene in botryococcenes extraction. On the other hand, p-cymene has been used to extract lipids from different biomass sources, including oleaginous microalgae (De Jesus and Maciel Filho, 2020). Even though we found a similar hydrocarbon extractability between p- cymene and ɣ-terpinene when the solvent content was 10% v/v (Fig. 3B), p-cymene was more aggressive on the cells than ɣ-terpinene (Fig. 3C). It has been previously reported that p-cymene was more toxic against insects than γ-terpinene, indicating the latter will likely be more appropriate for future botryococcene milking setups (Gong and Ren, 2020). The results indicate that ɣ-terpinene could be used as an alternative solvent to replace hexane for *in situ* extractions of hydrocarbons, but only for short extraction times. Different bioprocess designs to the *in situ* extraction procedure used in our study could help to reduce the toxic effects of solvent contact on algal cells. For example, it was recently shown that *B. braunii* cultures grown in a bubble column bioreactor attached to an extraction column could achieve a long-term *in situ* extraction of extracellular hydrocarbons over 30 days by using a daily extraction time of 5 h with hexane (Kleinert and Griehl 2022). Other *in situ* extraction systems and setups, such as membrane filtration, have shown to be successful in extracting exopolysaccharides (EPS) from *B. braunii* CCALA778 by using a microfiltration hollow-fiber membrane without requiring solvent (García-Cubero *et al.,* 2018). A similar system was used to continuously extract the heterologous sesquiterpenoid patchoulol with a hydrophobic hollow-fiber membrane acting as a physical interaction contact matrix between engineered algal cells and the solvent dodecane (Overmans *et al*., 2022).

Hexane is a suitable extraction solvent due to its low boiling point (69 °C), low cost, and high extraction efficiency (Jackson *et al*., 2017; Ennaceri *et al*., 2023). Using hexane facilitates solvent separation from the botryococcenes by distillation, as they have boiling points well above hexane (> 300 °C; Jackson *et al*., 2017). Due to its higher volatility, hexane would be superior to brominated alkanes and the terpene green solvents with boiling points > 150 °C, but only if the downstream process relies on energy-intensive distillation (Table 1). Other emerging techniques, including organic solvent nanofiltration and alternative wetting of membranes, may enable low-energy separations of botryococcenes, similar to those recently reported for sesquiterpene alcohols (Jackson *et al*., 2017; Alduraiei *et al*., 2021; Overmans *et al*., 2022). Terpene green solvents used in this work (134–136 g/mol; Table 1) have smaller molecular weights than botryococcenes (e.g., 466.8 g/mol C_34_H_58_), which would facilitate their downstream separation by these techniques. Although p-cymene may be the least expensive alternative solvent when it compares to hexane for botryococcene extraction (Table 1), solvent recycling of p-cymene and other green solvents would make it more competitive for the downstream processing of algae-based biofuels. Furthermore, the improved extraction efficiencies and sustainable nature of green solvents make them attractive alternatives to hexane.

## Conclusions

Using brackish cultivation to improve hydrocarbon extractability is a step-change in milking efficiency potential for *B. braunii* cultures, but the solvent choice was limited to the use of straight-chain alkanes. Here, we show that terpene green solvents and heavy solvents like bromodecane could also be used as alternative solvents to hexane for milking botryococcenes from *B. braunii* race B cultured in brackish medium. However, they are less biocompatible than dodecane, leading to 33–100% chlorophyll content loss. Our results warrant follow-up testing of these parameters in short-time scale solvent contactor systems that may improve the *B. braunii* hydrocarbon milking processes.

## Declarations

## Supporting information

Supplemental File 1 Data of all Figures

## Acknowledgments

We would like to thank Chandrasekaran Lakshmipathy of the KAUST Lab Equipment Maintenance (LEM) team for assistance in upgrading and initializing the GC-FID-MS unit.

## Authors’ contributions

TAO-U performed the experiments, contributed to the research planning, experimental design, data collection and visualization, formal analysis, and wrote the original draft manuscript. SO contributed to the research planning, experimental design, GC-MS analysis, and data visualization, and reviewed the manuscript. BBdF contributed to experimental design, assisted in the operation of Algem photobioreactors, and reviewed the manuscript. EL-G contributed to funding acquisition and data visualization, and reviewed the manuscript. The research was mainly conducted in the laboratory of KJL, who was responsible for experimental design, project scope, funding acquisition and reviewing the manuscript. All authors read and approved the final manuscript.

## Availability of data and materials

All data generated or analyzed during this study are included in this article and its supplementary information file.

## Ethics approval and consent to participate

Not applicable.

## Consent for publication

All the authors consented to the publication of this work.

